# Use of Transabdominal Ultrasound for the Detection of Intra-Peritoneal Tumor Engraftment and Growth in Mouse Xenografts of Epithelial Ovarian Cancer

**DOI:** 10.1101/2020.01.20.912402

**Authors:** Laura M. Chambers, Emily Esakov, Chad Braley, Mariam AlHilli, Chad Michener, Ofer Reizes

## Abstract

**Objective:** To evaluate intraperitoneal (IP) tumor engraftment, metastasis and growth in a pre-clinical murine epithelial ovarian cancer (EOC) model using both transabdominal ultrasound (TAUS) and bioluminescence *in vivo* imaging system (IVIS).

**Methods:** Ten female C57Bl/6J mice at six weeks of age were included in this study. Five mice underwent IP injection of 5×10^6^ ID8-luc cells (+ D-luciferin) and the remaining five mice underwent IP injection of ID8-VEGF cells. Monitoring of tumor growth and ascites was performed weekly starting at seven days post-injection until study endpoint. ID8-luc mice were monitored using both TAUS and IVIS, and ID8-VEGF mice underwent TAUS monitoring only. Individual tumor implant dimension and total tumor volume were calculated. Average luminescent intensity was calculated and reported per mouse abdomen. Tumor detection was confirmed by gross evaluation and histopathology. All data are presented as mean +/- standard deviation.

**Results:** Overall, tumors were successfully detected in all ten mice using TAUS and IVIS, and tumor detection correlated with terminal endpoint histology/ H&E staining. For TAUS, the smallest confirmed tumor measurements were at seven days post-injection with mean long axis of 2.23mm and mean tumor volume of 4.17mm^3^. However, IVIS imaging was able to detect tumor growth at 14 days post-injection.

**Conclusions:** TAUS is highly discriminatory for monitoring EOC in pre-clinical murine model, allowing for detection of tumor dimension as small as 2 mm and as early as seven days post-injection compared to IVUS. In addition, TAUS provides relevant information for ascites development and detection of multiple small metastatic tumor implants. TAUS provides an accurate and reliable method to detect and monitor IP EOC growth in mouse xenografts.

## Introduction

Epithelial ovarian cancer (EOC) is a leading cause of gynecologic cancer related mortality in women [1]. The five-year overall survival for women with EOC is poor since the majority of patients present with advanced and metastatic disease [2]. Additionally, although patients initially respond well to treatment with surgery and chemotherapy with carboplatin and paclitaxel, the vast majority of women will recur [4-8]. Ovarian carcinomas primarily undergo peritoneal dissemination, and are often associated with malignant ascites. This pattern of spread is associated with vague symptoms which leads to delays in diagnosis [3]. There is a significant unmet need for methods to facilitate early diagnosis of EOC and advance current therapeutic options.

Pre-clinical research utilizing EOC cell lines and patient-derived xenografts shows tremendous promise in advancing the current understanding of EOC carcinogenesis and therapeutics [9-15]. In longitudinal pre-clinical studies, the ability to detect tumor engraftment and sequentially assess tumor volume utilizing non-invasive techniques is essential to assessing tumor growth and treatment response. However, despite the existence of many cell lines that closely replicate human EOC at a cellular level, difficulty monitoring intraperitoneal tumor formation, growth and metastasis remains a major limitation in the execution of preclinical EOC studies [9-15].

In-vivo monitoring of EOC cell lines can be accomplished using several well-developed techniques including RFP, GFP, luciferase and ROSA reporter systems [16, 17]. Bioluminescence *in vivo* imaging system (IVIS) using luciferase reporter containing cell lines has been commonly utilized to track tumor growth over time, but this technique has limitations. This imaging technique involves injection of luciferin in conjunction with tumor cells, which is invasive and can initiate an inflammatory response [18]. Additionally, this technique only provides qualitative information regarding tumor progression [17]. The primary concern for use of IVIS is the necessity to use modified cell lines which have a tendency for genetic drift and phenotypic alterations. As such, use of IVIS for monitoring of patient-derived xenografts is not feasible [19,20]. In addition, a pertinent characteristic of EOC patients is the development of ascites throughout the progression of disease and the accuracy of luciferase is diminished in the presence of abdominal ascites as a result of dilution [21-24].

In clinical practice, transabdominal ultrasound (TAUS) is frequently utilized in the evaluation of women with gynecologic diseases, including EOC [25,26]. Despite being non-invasive, cost-effective and accurate, data for use of TAUS for monitoring of EOC in pre-clinical murine models is limited [27]. The objective of this study was to evaluate intraperitoneal tumor engraftment and growth in the presence and absence of ascites in a pre-clinical murine model of EOC utilizing both TAUS and IVIS imaging.

## Methods

### Cell Lines and Lentiviral Transformation of ID8 Cells with Luciferase Vector

ID8 and ID8-VEGF syngeneic EOC cell lines were cultured in Dulbecco Modified Eagle Medium (DMEM) media containing heat inactivated 5% FBS (Atlas Biologicals Cat # F-0500-D, Lot F31E18D1) and 100 U/mL penicillin-streptomycin and 1% insulin/transferrin/selenium and grown under standard conditions. HEK 293T/17 (ATCC CRL-11268) cells were plated at 65 % confluence in a 100 mm dish and cultured in 9 mL DMEM supplemented with heat inactivated 10% FBS (Atlas Biologicals Cat # F-0500-D, Lot F31E18D1)[15]. ID8 cells were subsequently transfected with luciferase containing construct pHIV-Luciferase #21375 4.5 µg (Addgene) to generate the ID8-luc cells. Briefly, 3 mL of the DMEM media was removed and ID8 cells were co-transfected with Lipofectamine 3000 (L3000015 Invitrogen) 35 µL of Plus reagent / 41 µL of Lipofectamine 3000, 3rd generation packaging vectors pRSV-REV #12253 4.3 µg, pMDG.2 #12259 4.3 µg, and pMDLg/pRRE #12251 4.3 µg (Addgene) and lentiviral vector directing expression of luciferase reporter pHIV-Luciferase #21375 4.5 µg (Addgene) in 3 mL of OptiMEM media. Following 8 hours of incubation, media of the 293T/T17 cultures was replaced and following 18 hours of incubation media containing viral particles were harvested and filtered through a 0.45 µm Durapore PVDF Membrane (Millipore SE1M003M00). Viral transfections were carried out over 72 hours ID8 parental cells and transduced cells were selected by their resistance to 2 μg/mL puromycin (MP Biomedicals 0219453910). Prior to use in this experiment, activity of luciferase promoter and tumor growth was confirmed in a pilot cohort of mice.

### Study Approvals

All animal work throughout the study was completed and approved by the Institutional Animal Care and Use Committee (IACUC) (Protocol #2018-2003) of the Biological Resource Unit of The Cleveland Clinic Foundation Lerner Research Institute. Post tumor cell injection, mice were monitored weekly for signs of distress and humane endpoint was reached upon development of tumor burden >150mm^3^ (by ultrasound) or debilitating ascites development as outlined in the above protocol, mice also reached humane endpoint if ruffled fur, reduced mobility, or hunched body posture was observed. Upon reaching humane endpoint criteria, mice were immediately euthanized by CO2 asphyxiation and cervical dislocation. No animals were found dead before meeting endpoint criteria in this study. All researchers participating in animal studies were appropriately trained by veterinary technicians or skilled lab personnel following approved IACUC guidelines.

The surgical specimens used to generate the PDX model were obtained with permission from the Institutional Review Board of the Cleveland Clinic Foundation under IRB#18-062 Gynecologic Oncology Tissue Collection. Per IRB#18-062 all specimens were collected after written informed consent was obtained from the patient. Tissue collected was frozen and stored and when available, live tissue procured to establish patient derived xenografts.

### Mouse Xenografts

Ten female C57Bl/6 mice were purchased from Jackson Laboratories (Bar Harbor, ME) at 6 weeks of age. After two weeks of acclimation, mice underwent IP injection of 300uL of either 5×10^6^ ID8-luc (n=5) or ID8-VEGF (n=5) cells. All mice met endpoint criteria by 58 days post cell injection and no mice were removed from the study prior to meeting endpoint criteria.

### Tumor Monitoring

Ultrasonography was performed using a Vevo2100 (VisualSonics) with an abdominal imaging package and MS550D probe (40Hz). TAUS surveillance was initiated seven days following IP tumor injection. TAUS was performed every seven days until study endpoint at 67 days. Upon TAUS imaging, mice were also monitored for signs of obvious physical distress as outlined in the approved IACUC protocol. Mice were anesthetized using isoflurane (DRE Veterinary) and placed in the supine position. Following the removal of abdominal hair using Nair (Church & Dwight Co. Inc.), sterile ultrasound gel was applied to the abdomen. TAUS was performed using Vevo2100 (VisualSonics) using the abdominal imaging package and MS550D probe (40Hz) (Figure 5 - Supplemental). For each mouse, the abdomen was assessed for tumor in four quadrants. Tumors were noted to be absent or present at each assessment. Tumor dimensions (length and width) were recorded and tumor volume was calculated using the formula: (Length*(Width^2^))/2.

### 2D IVIS imaging

Bioluminescence images were taken within 48 hours of ultrasound images with IVIS Lumina (PerkinElmer) using D-luciferin as previously described [24]. Mice received an IP injection of D-luciferin (Goldbio LUCK-1G, 150mg/kg in 150mL) under inhaled isoflurane anesthesia. Images were normalized (Living Image Software) with a minimum and maximum radiance of 7.5810^5^ and 5.3910^8^ photons/second/cm^2^/steradian, respectively. All images were obtained with a 15 second exposure. Average luminescent intensity in photons per second/cm^2^/steradian was calculated and reported for each mouse abdomen.

### 3D IVIS Imaging

Upon endpoint, bioluminescence and x-ray images were taken using the IVIS Spectrum system (PerkinElmer). Mice were sedated with 2% isoflurane (DRE Veterinary) inhalation in an airtight transparent anesthesia box for 5 minutes. Mice were shaved front and back and Nair was applied to remove remainder of the hair before being IP injected with D-luciferin (Goldbio LUCK-1G, 150mg/kg in 150 mL). Mice are placed in a supine position on the light-tight chamber of the CCD camera imaging unit. Sequential images were acquired at 1min intervals (60 s exposure, no time delay) for at least 30 min. The luminescence camera was set to 60 s exposure, medium binning, f/1, blocked excitation filter, and open emission filter. The photographic camera was set to auto exposure, medium binning, and f/8. Average luminescent intensity in photons per second/cm^2^/steradian was calculated and reported for each mouse abdomen. Identical settings were used to acquire each image and region of interest during the study. Ultrasound and IVIS imaging were performed independently by two separate investigators who were blinded to the results of the other imaging modality.

### Statistics

All data are presented as mean +/- standard deviation. Tumor volumes presented as mean+SEM and graphed over time. All statistical analysis was performed in GraphPad Prism v8.

## Results

Tumor engraftment was detected in all C57Bl/6J via TAUS between 7-14 days, gross examination at necropsy and on histopathology. In addition, in mice injected with ID8-luc, tumor engraftment was noted at 14 days. In all cases, EOC tumors were detected before any clinical signs (ascites, palpable masses, lethargy). Beginning at Day 7 post-injection, TAUS was performed and a maximum of four tumor measurements were recorded per mouse in each abdominal quadrant. Six mice (60%) had one detectable tumor on TAUS at 7 days. All mice (n=10) had at least one tumor detectable on TAUS at 14 days post-injection and 20% (n=2) had two detectable tumors. Mean tumor dimensions and volumes for ID8-luc and ID8-VEGF are displayed in Table 1 and Table 2, respectively.

**Table 1.**

|  | Transabdominal Ultrasound Measurement |  |  |  | IVIS Imaging |  |
| --- | --- | --- | --- | --- | --- | --- |
|  | Mean Tumor Implant Long Axis (mm) (n=20) | Mean Tumor Implant Short Axis (mm) (n=20) | Mean Tumor Implant Volume (mm3) (n=20) | Mean Total Tumor Burden per Mouse (n=5) (mm3) |  | photon/second/cm^3/sr |
| Day 7 | 2.83 | 1.80 | 4.61 | 4.61 | Day 7 | 2.63e6 |
| Day 14 | 2.92 | 2.11 | 6.81 | 9.55 | Day 14 | 1.11e6 |
| Day 21 | 4.10 | 2.58 | 14.5 | 43.38 | Day 24 | 4.15e6 |
| Day 28 | 4.26 | 2.76 | 17.28 | 69.10 | Day 31 | 4.88e6 |
| Day 35 | 4.94 | 3.50 | 32.54 | 130.14 | Day 39 | 7.75e6 |
| Day 42 | 5.88 | 4.14 | 51.34 | 205.35 | Day 46 | 1.95e<sup>7</sup> |
| Day 49 | 7.20 | 4.58 | 75.58 | 305.53 | Day 57 | 2.18e<sup>7</sup> |
| Day 56 | 8.56 | 5.06 | 110.00 | 432.83 |  |  |

**Table 2.**

|  | Transabdominal Ultrasound Measurement |
| --- | --- |

|  | Mean Tumor Implant Long Axis (mm) (n=20) | Mean Tumor Implant Short Axis (mm) (n=20) | Mean Tumor Implant Volume (mm<sup>3</sup>) (n=20) | Mean Total Tumor Burden per Mouse (n=5) (mm<sup>3</sup>) | Presence of Ascites |
| --- | --- | --- | --- | --- | --- |
| Day 7 | 2.23 | 1.98 | 4.39 | 4.39 | No |
| Day 14 | 3.00 | 1.90 | 5.56 | 6.79 | No |
| Day 21 | 3.98 | 2.54 | 13.52 | 35.23 | Yes |
| Day 28 | 4.30 | 2.70 | 16.73 | 49.92 | Yes |
| Day 35 | 5.36 | 3.70 | 38.59 | 130.28 | Yes |
| Day 42 | 6.55 | 4.03 | 56.09 | 178.81 | Yes |

The smallest tumor short and long axis measurements detected at 7 days were 1.74mm and 2.23mm, respectively. The lowest recorded tumor volume was 4.17mm. Ascites was detected as early as 21 days. Tumor volume detected via TAUS over time (method described in Supplemental Figure 1) is displayed for both ID8 and ID8 VEGF mice in Figure 1. Figure 2 depicts weekly TAUS images of EOC tumor implant over time in ID8-luc without ascites (A) and in ID8-VEGF (B).

**Figure 1.**
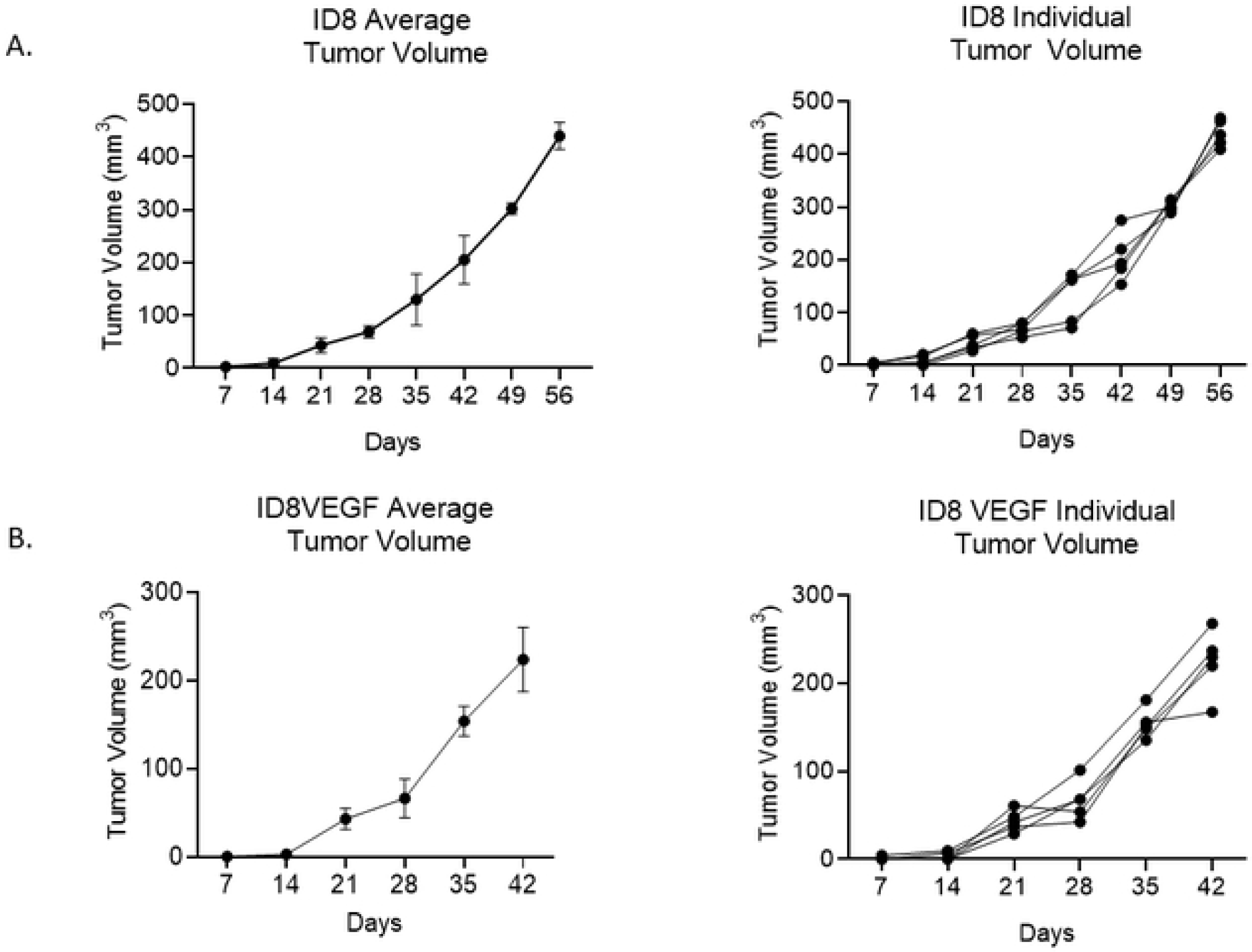
TAUS Allows for Monitoring of Tumor Engraftment and Growth in Mice with Ovarian Cancer Xenografts of ID8 (A) and ID8-VEGF (B). Figures demonstrate average total tumor volume and weekly total tumor volume per individual animal (n=5).

**Figure 2.**
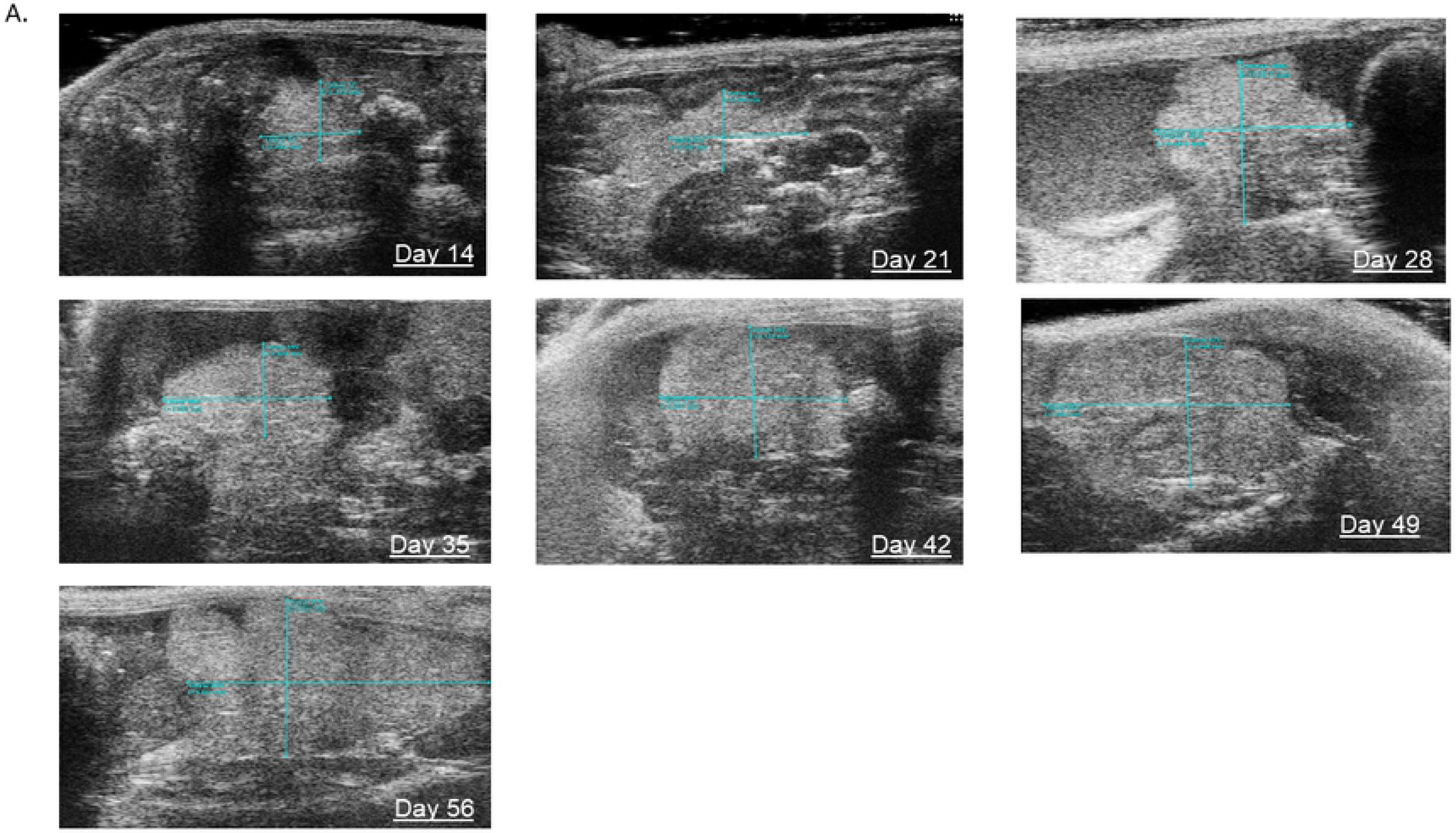
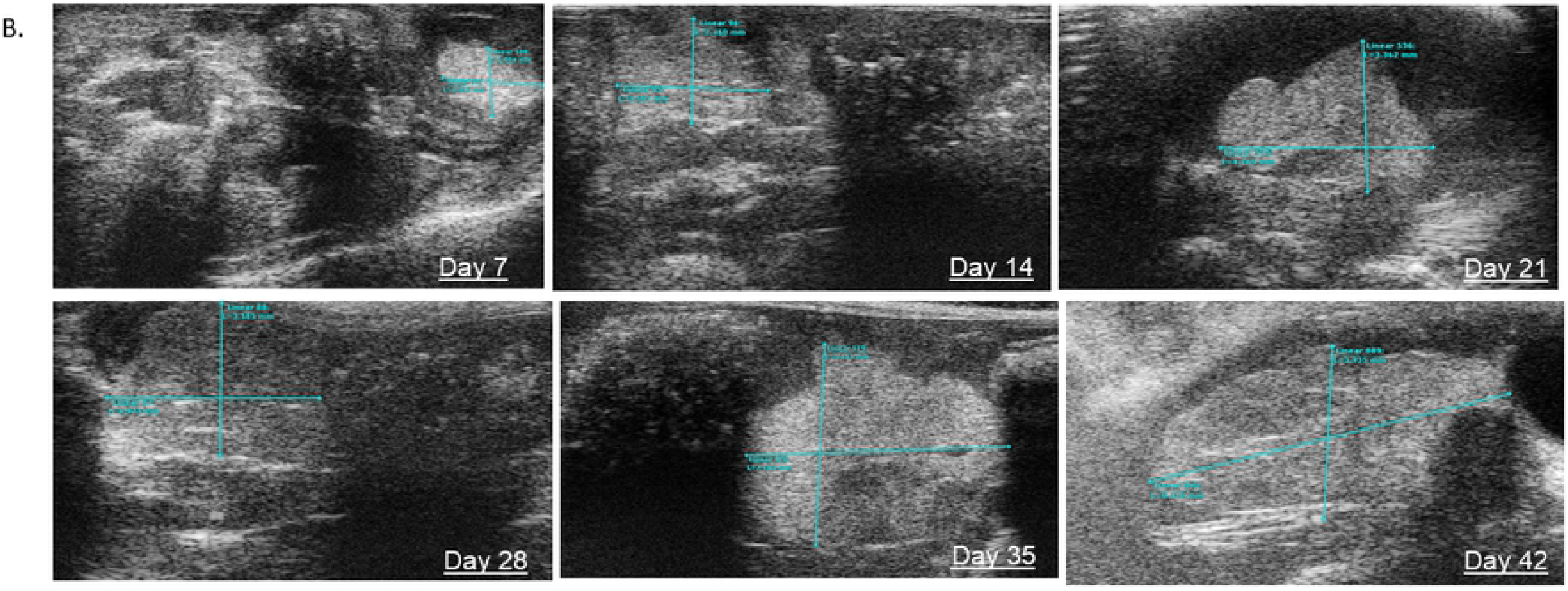
Transabdominal ultrasound demonstrating ability to monitor tumor implant .longitudinally in ID8-Luc (A) and ID8-VEGF (B) mice. Cyan colored caliper measurements can be observed at each time point which was utilized to monitor tumor volume.

Supplemental Figure 1. Procedural Steps for Transabdominal Ultrasound. Once the imaging unit is initialized, the warming plate and heart monitor are turned on (A). After induction of anesthesia with inhaled isoflurane, mice are placed in a supine position on the platform to monitor heart rate (B), and the abdominal fur is removed (C). Sterile ultrasound gel is then applied to the abdomen and TAUS performed using Vevo2100 using the abdominal imaging package and MS550D probe (40Hz) (D). Once a tumor is identified (E), the image is captured and length and width are measured (F).

Within 48 hours of TAUS, 2D IVIS imaging was performed. Tumor detection by 2D IVIS imaging was noted at 14 days post cell injection and intraperitoneal tumor growth over time was tracked as previously reported (Figure 3A, B). Additionally, 3D IVIS imaging including murine x-ray was performed at endpoint to determine tumor location (Figure 3C).

**Figure 3.**
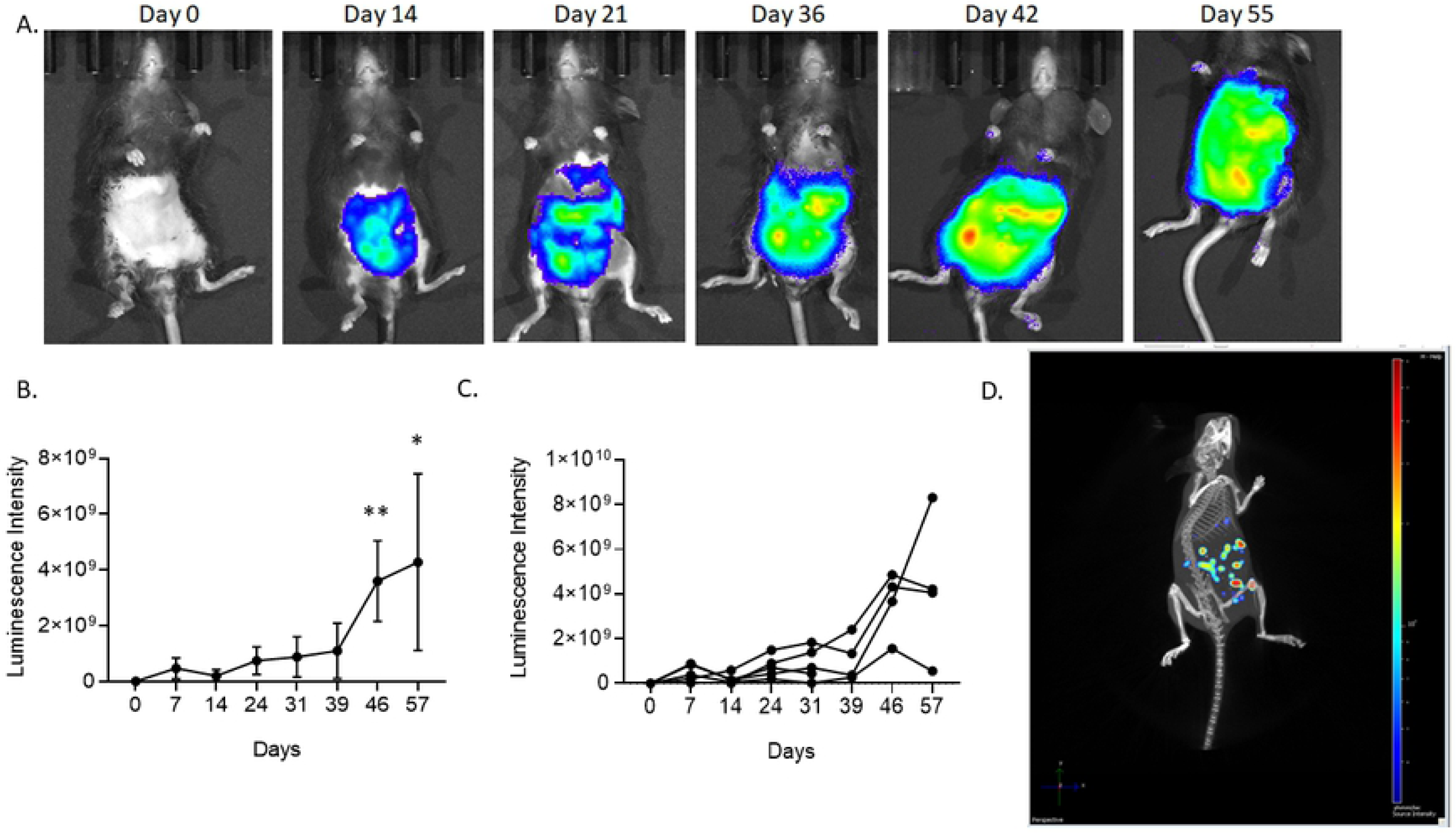
2D IVIS imaging tracked tumor growth over time, and 3D IVIS imaging determined endpoint tumor volume in ID8 tumor bearing C57Bl/6 mice. A. A representative time course of 2D IVIS imaging to track ID8 growth in C57Bl/6 mice. B. Graph depicting an average of ID8 tumor growth over time by fluorescence intensity (n=5). C. Graph depicting individual ID8 tumor growth over time. D. An endpoint 3D IVIS depiction of tumor volume and location. *p<0.05, **p<0.01

Prior to necropsy, the murine abdominal cavity was imaged to confirm gross tumor presence in the ID8 and ID8 VEGF cohort. Each tumor was then excised and stained using H&E to confirm EOC histology (Figure 4). Following murine necropsy, ID8 tumors were imaged and excised (Figure 4). ID8 and ID8 VEGF EOC tumor phenotype was confirmed by histology (Figure 4A and B respectively).

**Figure 4.**
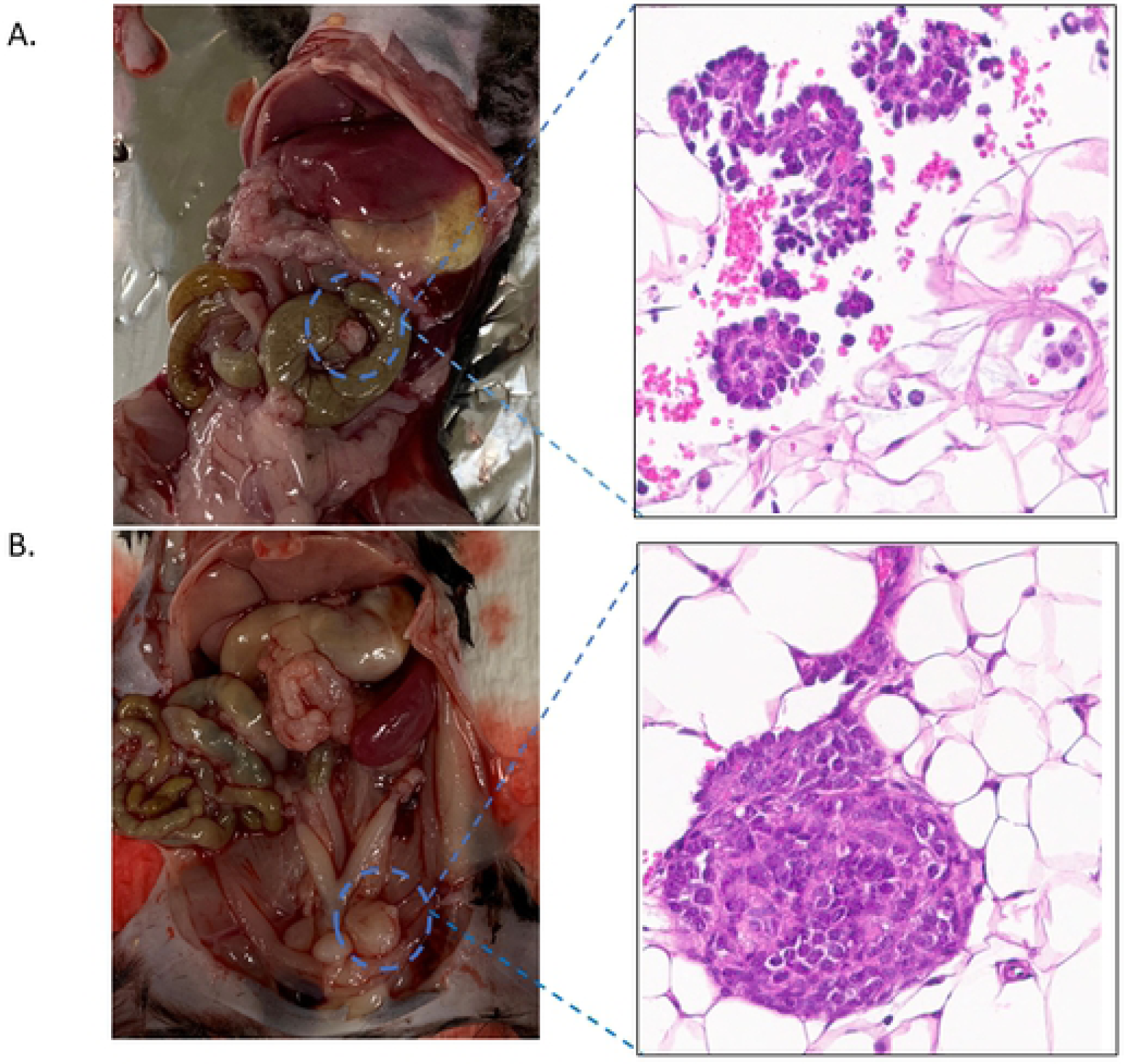
Upon necropsy macroscopic and histologic EOC tumors were identified and validated for both ID8 and ID8 VEGF cell lines. A. Murine necropsy showing ID8 tumor mass (blue dotted circle), and resulting H&E stain confirming EOC pathology. B. Murine necropsy showing ID8 VEGF tumor mass (blue dotted circle), and resulting H&E stain confirming EOC pathology.

Additionally, to test whether TAUS can be used to detect PDX tumors, we injected mice with a PDX single cell suspension of EOC human cells. We were able to detect tumor growth at 10 days post injection (Figure 5).

**Figure 5.**
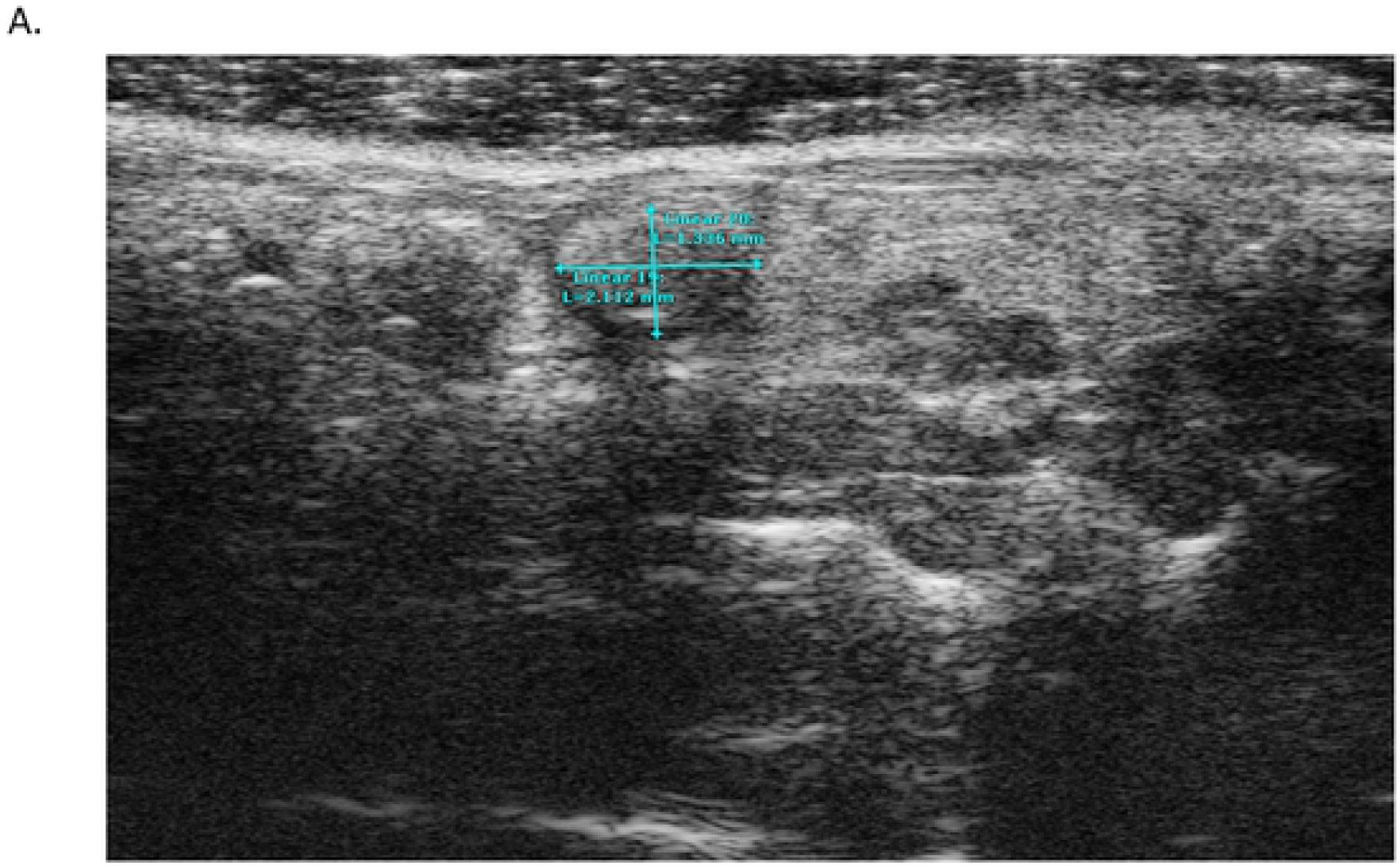
Human derived EOC PDX tumor detected at 10 days post IP injection (A).

## Discussion

Mouse xenografts represent an important method to pursue urgently needed preclinical studies to understand pathogenesis and develop new therapies for EOC. While many orthotopic models exist that closely mirror human EOC, techniques to monitor intraperitoneal tumors in an accurate, non-invasive fashion are limited. In this study, we applied ultrasonography, to evaluate the engraftment and growth of EOC in a pre-clinical model. We demonstrated that in murine models of EOC, TAUS can be used to accurately detect and monitor the growth of EOC xenografts with tumors and ascites detected as early as 7 and 21 days post-injection, respectively. We found TAUS is more sensitive for detection of disease progression compared to bioluminescence assays where tumor detection first occurred at 14 days post-injection.

Currently utilized and previously described strategies for tumor monitoring in murine models of EOC fall short [9-15, 21-24]. IVIS imaging is frequently used for tumor assessment in murine models of EOC but has significant limitations. First, the cell-line must contain a luciferase reporter, which limits the ability to utilize high fidelity patient-derived tumor graft models. Second, concerns exist regarding initiation of an inflammatory response or other phenotypic and genotypic alterations that may render the cell line less applicable to human EOC [19-21, 23]. Finally, detection of ascites is compromised in IVIS models. Baert et al demonstrated that reduced sensitivity of IVIS in the presence of luciferase with a significantly decreased in the presence of ascites within an ID8-luc model [23]. As the majority of human and mouse EOC lines have a penchant for ascites development, the detection of ascites is of high importance. In the clinical setting, ascites significantly impacts patient quality of life and is a harbinger of advanced, progressive disease. Ascites is important to study in pre-clinical translational models as it can yield diagnostic and prognostic information.

In clinical practice, TAUS is frequently utilized in the evaluation of women with gynecologic diseases, including EOC [25, 26]. However, prior to this study, application of ultrasonography to murine pre-clinical EOC models has been limited. Weroha et al. utilized ultrasonography to assess tumor growth in patient-derived xenografts of EOC with high correlation between ultrasound assessment and tumor measurements at necropsy [27]. In addition, TAUS has been utilized in pre-clinical models of non-gynecologic intra-abdominal cancers, including pancreatic and genitourinary malignancies [28-30]. Within a murine model of bladder cancer, Patel et al. demonstrated high correlation between tumor size with transabdominal micro-ultrasound and at necropsy and were able to detect tumors as small as 0.95 mm^3^ [30]. Similarly, in pre-clinical murine models of pancreatic adenocarcinoma, intra-pancreatic tumors were detected as early as three days post-injection, and tumor metastasis in addition to ascites was identified in all animals at two weeks with excellent correlation between tumor volume and [29].

TAUS offers several potential advantages over currently available imaging tools for the monitoring of murine models of EOC. Primarily, we demonstrate in this study that tumor detection can be assessed as early as one week post-injection, with tumor implants detected as small as 2mm in longest dimension. Secondly, malignant ascites and innumerable tumor implants are pathognomonic of human EOC. This method allows researchers to monitor treatment response via tumor volume and ascites in parallel to patients undergoing chemotherapy where radiologic scoring systems such as RECIST criteria are used. In addition, TAUS can be utilized for EOC monitoring in cell lines that do not have RFP, GFP, luciferase or ROSA reporter systems. Thereby, this allows for in-vivo monitoring of any intra-peritoneal EOC cell line with or without ascites development, including PDX models. This is important because it allows researchers to follow tumor growth and treatment response over time with cells transplanted directly from patient tumor specimens without the need for luciferase transduction. Finally, the ability to accurately detect tumors may represent a strategy to minimize animal euthanasia, as their disease burden can be monitored in-vivo to end-point during an experiment without need for early necropsy with each animal serving as its own control. Therefore, monitoring EOC growth and response via TAUS has improved detection, higher sensitivity and increased breadth and utility over presently utilized imaging techniques.

In clinical practice, transabdominal and transvaginal US remain gold-standard for the initial assessment of gynecologic pathology, including ovarian tumors. TAUS is non-invasive and cost-effective with low risk to the patient. In this study, we demonstrate that this same imaging modality can be applied to mouse xenografts. Based on these results, we have adopted TAUS as a method to monitor tumor growth and treatment response in EOC preclinical studies in both syngeneic and PDX models with excellent success and reproducibility. One limitation of this model is the need for mouse anesthesia during TAUS. In this series, anesthesia and TAUS were well tolerated by the mice with no adverse intra-anesthesia events or mortalities related to the procedure. Despite this, to the best of our knowledge, this study represents the first publication assessing the feasibility of TAUS for preclinical murine models of EOC in parallel with IVIS imaging.

In conclusion, TAUS shows promise in the detection of tumor growth and metastasis and response to therapies in intraperitoneal mouse xenografts of EOC. TAUS allows for detailed measurements of tumors and metastatic implants, ascites and is more sensitive than IVIS imaging.

## Author Contributions

Chambers and Reizes were responsible for the conception and planning of the study. Chambers, Esakov, Braley, Michener, AlHilli and Reizes participated in study design. Chambers, Esakov, Braley performed data collection and management. Chambers, Esakov and Reizes designed and performed the data analysis. Chambers and Esakov drafted the manuscript. All authors critically revised the manuscript and all authors approved the manuscript in its final version for publication.

